# The dietary polyphenol metabolite protocatechuic acid produces acute and sustained effects on hippocampal electrical activity

**DOI:** 10.1101/2023.01.12.523723

**Authors:** Marta Montero-Atalaya, Ricardo Muñoz-Arnaiz, Julia Makarova, Begoña Bartolomé, María-Victoria Moreno-Arribas, Oscar Herreras

## Abstract

Dietary polyphenols and in particular bioavailable metabolites resulting from gut microbiota transformations appear to have beneficial effects in situations of impaired cognition, combatting memory deficits in acute pathological models of neurodegeneration. Modifications to blood flow may underlie the effects of these molecules and although some such metabolites cross the blood-brain barrier, their targets and electrophysiological effects remain unknown. Hence, we explored the systemic and direct effects of protochatechuic acid (PCA) on electrical activity in the hippocampus and cortex of anesthetized female rats, recording evoked and spontaneous high-density field potentials (FPs) to mathematically derive pathway-specific FP generators. We found transient and sustained effects of PCA on evoked activity in the CA1 field, including paradoxical actions on excitatory transmission that depend on the route of administration. Systemic delivery of PCA altered the ongoing activity of some FP generators, albeit with marked inter-animal variation. Interestingly, PCA induced the detachment of infraslow cortico-hippocampal activities over a scale of minutes. These results point to direct actions of polyphenols on cell and network electrical activity, some of which reflect non-specific actions. Thus, dietary-derived polyphenols appear to fulfill neuromodulatory roles, encouraging the search for additional targets to better guide their use in preventing brain pathologies.

## Introduction

There is evidence that a polyphenol-rich diet plays an important role in the prevention of neurodegenerative diseases, cardiovascular accidents and stroke. It also appears to improve the cognitive deficits associated with aging and those related to malnutrition in children.^1-5^ These phenomena have been at least partially associated with the vascular effects of these substances, which include angiogenesis or increased blood flow.^6,7^ A number of effects on signaling pathways have been reported at the cellular level,^8,9^ yet mechanistic data is on the whole scarce, particularly regarding the electrophysiological effects of polyphenols on specific brain structures.

Dietary polyphenols exert some effects by interacting with the gut microbiota, with intestinal bacteria metabolizing them into specific metabolites that become bioavailable. As a result of these biotransformations in the gastrointestinal tract, novel metabolites enter circulation that are not present in food and that may exist in only minor quantities.^10-12^ Protocatechuic acid (PCA, or 3,4-dihydroxybenzoic acid) is one of the main metabolites of complex polyphenolic compounds like anthocyanins and flavan-3-ols, phenols that have been widely studied for their beneficial impact on human health through their anti-inflammatory, antitumoral, immunoregulatory and neuroprotective effects.^13,14^ Moreover, several studies have demonstrated the ability of gut microbial-derived metabolites to cross the blood–brain barrier (BBB)^15^, suggesting that they might be responsible for the protective effects of dietary polyphenols on brain function and cognitive performance.^16-17^ Thus, in addition to possible vascular actions, it has been suggested that polyphenols and their metabolites may directly modulate brain activity.^18-19^

There have been a few studies into the electrophysiological effects of polyphenols and in general, they have focused on very specific phenomena like long-term potentiation.^20^ Given the slow temporal dynamics of the cognitive and pathological events in which polyphenols have been implicated, it would be particularly relevant to explore any sustained and long-term changes to electrical activity. Indeed, such alterations have been shown to play an important role in some neurodegenerative diseases like Parkinson’s and Alzheimer’s disease^21,22^. Therefore, the main objective of this study was to explore the effects of PCA on electrophysiological activity in the hippocampus and cortex of anesthetized adult rats. These structures were chosen as they are critically involved in memory processes, and they are also significantly affected by neurovascular accidents and neurodegenerative diseases in which polyphenols have been shown to have a favorable effect.^18,19^ In addition, we complemented this study by exploring the effects of PCA on hippocampal-dependent behavioral tasks.

Since PCA is known to have dose-dependent antagonistic effects (oxidant-antioxidant), it is essential to consider the pharmacokinetics and toxicity of this compound.^14^ Thus, we tested these features of PCA by (1) direct delivery into the brain parenchyma and (2) through systemic intraperitoneal delivery enabling this compound to be transported to and metabolized in the brain. The effects on membrane and synaptic events were explored by studying hippocampal evoked potentials in the CA1 region. The exploration of systemic variables on brain network activity benefits from studying spontaneous field potentials (FPs). Hence, we used high density linear arrays to record across the cortex and hippocampus, employing spatial discrimination techniques to obtain the FP generators with independent dynamics, such as independent component analysis (ICA).^23^ This approach provides information on connectivity, since the generators have been formerly characterized in these structures and most are pathway-specific.^24,25^ The time course of the separated FP generators is also more convenient for quantitative studies over long periods, since they display varying electrophysiological patterns over different time scales.^26^

The results obtained indicate that PCA produces an effective and direct acute modulation of electrophysiological activity in the brain. We found sustained and transient effects of PCA that included paradoxical effects on hippocampal excitatory transmission depending on the route of administration. Interestingly, we found PCA induced detachment of cortico-hippocampal coherence in the rat.

## Methods

All experiments were performed in accordance with EU (2010/63/UE), Spanish (RD 53/2013) and local (Autonomous Community of Madrid, Order 4/8/1988) regulations regarding the use of laboratory animals, and the experimental protocols were approved by the Research Committee of the Cajal Institute. Twenty-six adult female Sprague-Dawley rats (250-300g) of between 3 and 4 months were used. Animals were inbred at the local animal facilities in a 12-hour light/dark cycle, stable temperature (20-22º C), and food (Teklad 2018) and water were *given ad libitum*.

We followed the ARRIVE Essential 10 guidelines to report on animal research. The experimental unit is an individual animal. Individuals were selected from different litters and randomly assigned to the different groups (n = 6 individuals per group). Sample size was determined according to former studies exploring the number of pathway-specific FP generators in the hippocampus.^25^ The activity within these generators varies over time and cannot be homogenized across animals,^25,26^ hence it cannot be used to determine sample size. Data analysis was blindly performed by researchers other than those who executed procedures.

### Experimental paradigm

Experiments were designed to explore the effects of directly applying PCA (Sigma Aldrich, ref. P-5630, CAS number 99-50-3) to the brain (15 mg/kg dissolved in artificial cerebrospinal fluid –ACSF-with 1% dimethyl sulfoxide -DMSO) or by systemic administration (i.p., 150 mg/kg dissolved in NaCl 0.9% with DMSO 1%: see Figure 1). These concentrations were selected based on existing evidence of brain uptake and of the plasma levels of phenolic metabolites generated following the ingestion of a polyphenol food source.^12,18,27^

**Figure 1.**
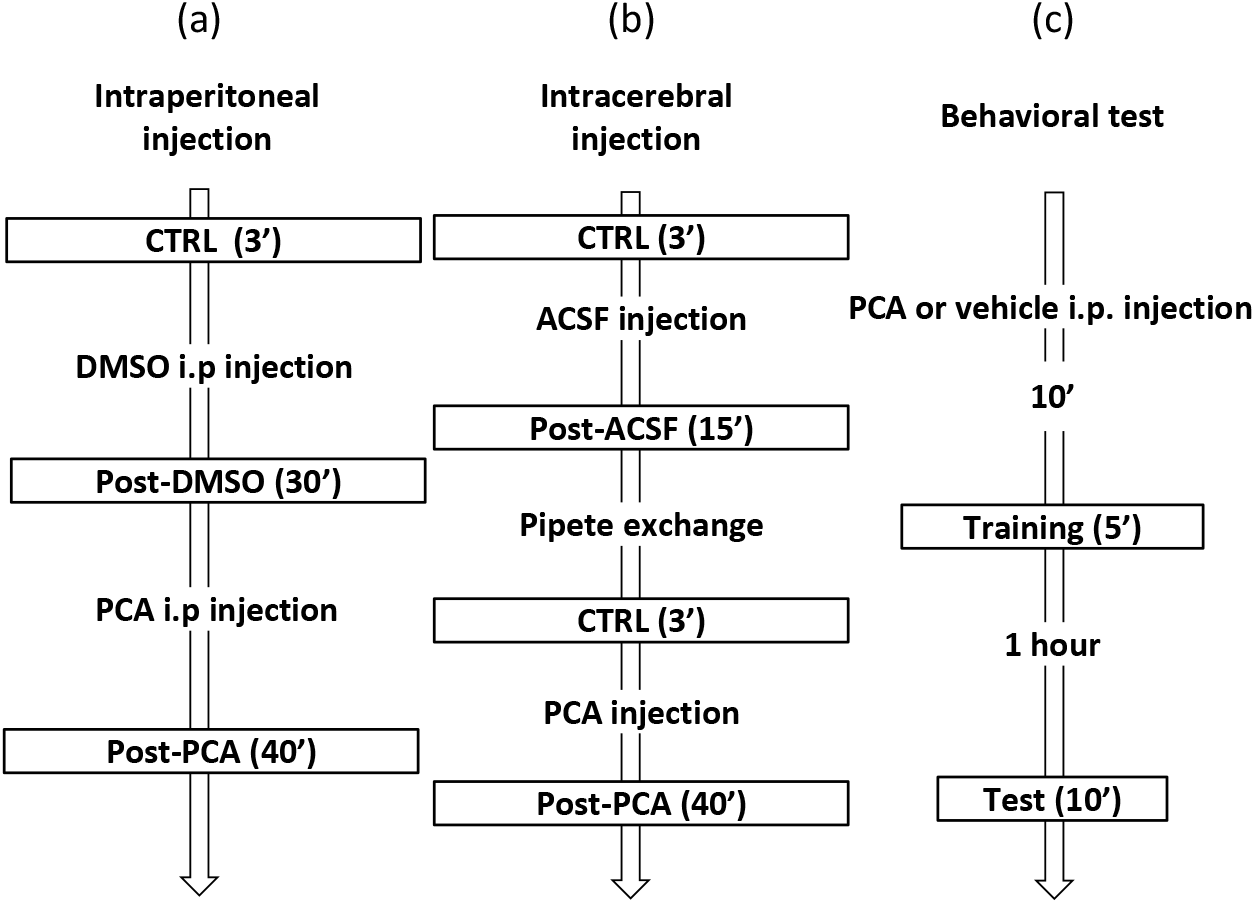
Experimental design and timeline of protocols. (a) The systemic effects of PCA were studied after administration via i.p., assessing the long-lasting or sustained effects on evoked potentials and spontaneous FP activity in the hippocampus and cortex as recorded through a pipette or multisite linear probe, respectively. Electrical activity was tracked for the periods marked in boxes. (b) The direct action of PCA on different membrane domains of neurons was investigated by microinjection through pipettes placed in the different layers of the CA1 region and studying layer-specific evoked potentials. Stimuli were delivered to the CA3 region (orthodromic) or the alveus (antidromic). The recording/injecting pipette filled with control solution was substituted for another filled with drug solution and placed 0.3-mm away to target fresh tissue. (c) Behavior was tested with the novel object recognition (NOR) and object location (OL) paradigms, applying them to the same animals 48 hours apart.

### Electrophysiological experiments

Experiments were performed in the dorsal hippocampus and the overlying cortex of individual animals anesthetized with urethane (1.5 g/kg, i.p.) and fastened to a stereotaxic device, maintaining their body temperature at 37 ± 0.1 °C. Surgical and stereotaxic procedures were performed as described previously.^28^ Concentric stimulating electrodes (TM33CCNON, WPI) were placed in the CA3b soma layer (in mm from bregma: AP -3.2; LM -2.6; V -3.3) to activate the Schaffer inputs to the CA1, or in the alveus (AP -5.5; LM -2.6; V -1.8) to antidromically activate the same population. Stimuli (0.07-0.1 ms square pulses) were delivered at 0.1 Hz. To explore the acute effects of PCA, glass micropipettes (5–10 MΩ) were filled with ACSF solution or supplemented with additional drugs (see below), and placed either at the st. pyramidale (anti and orthodromic population spikes: a-PS, o-PS) or at the st. radiatum (field excitatory postsynaptic potentials, fEPSPs).^29,30^ Microdrops (Ø∼50 μm) of the drug solutions or of the vehicle alone (DMSO 1%) were delivered by pressure microejection (all chemicals were purchased from Sigma-Aldrich). Spontaneous laminar FP activity was recorded through one multisite linear probe (32 sites, 100 μm: from Atlas, Leuven, Belgium or Neuronexus, Ann Arbor, MI) that spanned the lower layers of the cortex, as well as the CA1 and Dentate Gyrus (DG) hippocampal subfields. A subcutaneous Ag/AgCl wire under the skin of the neck was used as reference, and the signals were amplified and acquired using a MultiChannel System (Reutlingen, Germany) or Open Ephys hardware and software (20 kHz sampling rate).

Wide-band FPs (0.1 Hz – 5 KHz) were recorded in consecutive 3 min periods and we focused on irregular hippocampal activity, which is the dominant state at the anesthetic dosage used here.^31^ When the animal displayed frequent theta episodes before drug testing, anesthesia was supplemented (a 10-20 % increase of the initial dose) and the animal was allowed to stabilize for 10 min. The frequency bands were defined as: delta (δ), < 3 Hz; theta (θ), 3 to 8 Hz; alpha (α), 8-16 Hz; beta (β), 16-25 Hz; gamma (γ), 30 to 80 Hz.

### Spatial discrimination of pathway-specific components in FPs

In the brain, spontaneous FPs are driven by multiple co-activating synaptic sources whose potentials spread and mix in the volume, provoking mutual distortion.^32^ We used the ICA to separate these components,^24,33^ each of which has a unique spatial profile that enables its identification and anatomical matching^34^ (see Ref. 25 for applications and restrictions). In this study we employed the kernel density ICA algorithm (KDICA)^35^ that is usually implemented in MATLAB. Briefly, the *u*_*m*_(*t*) signals recorded are considered as the weighted sum of the activities of *N* neuronal sources or FP generators:

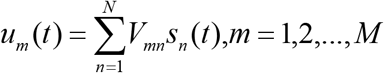

where (*V*_*mn*_) is the mixing matrix derived from the so-called voltage loadings or spatial weights of *N* FP generators on *M* electrodes, and *s*_*n*_(*t*) is the time course of the *n-th* FP generator. Using *u*_*m*_(*t*), the ICA finds both (*V*_*mn*_) and *s*_*n*_(*t*). Experimentally and through realistic computer models, it has been shown previously that the time-course of a FP-generator *s*_*n*_(*t*) corresponds to a postsynaptic temporal convolution of spike output in a population afferent to that recorded^36,37^. Dimension reduction was performed with a principal component analysis (PCA) prior to the ICA. Once extracted, each FP generator can be analyzed independently in the time or frequency domains. Typically, maintaining the 6-8 main PCA components optimizes ICA performance in recordings that span the dorsal CA1 and the DG.^38^ We routinely discarded noisy components with a total compound variance below 1% (i.e. always keeping 99% of the original FP variance). The time evolution of the power of an FP generator (in mV^2^) was calculated as indicated previously^39^ and the power of the FP components was estimated at the site where maximal amplitude was found.

### Behavioral experiments

Two behavioral tests were performed to evaluate the effect of systemic PCA on hippocampal-dependent memory (Figure 1), the novel object recognition (NOR) and object location (OL) tests.^40^ The NOR test evaluates an animal’s ability to recognize a novel object in a given environment without any reinforcement from external cues. We used a two-phase protocol, each with a 5 min habituation period. In an initial (training) phase, the animal was placed in the center of the box (40×40×40 cm) containing two objects (A and B) and it was allowed to explore them for 5 minutes. One hour later the test phase was carried out, in which one of the objects was changed (A and C) and the animal was allowed to explore for 10 minutes. The OL test evaluates memory with a spatial component and it follows a similar protocol to the NOR, except that in the test phase the same objects are used (A and B) but one of them has been moved from its initial position. In both paradigms, a single i.p. dose of PCA (150 mg/kg) diluted in saline with 1% DMSO (PCA group) was delivered 10 minutes before the test began, which was compared with a sham group that were administered only saline with 1% DMSO. A discrimination index was calculated from the time spent exploring the novel and the known objects relative to the total exploration time.

### Statistical analysis

The normality of the data was assessed using the Shapiro-Wilk test and the homogeneity of variances was checked with the Levene test. Differences between the groups were analyzed using a t-test for independent samples with a normal distribution; otherwise the non-parametric Mann Whitney U test was employed. The experiments involving i.p. injections had 3 levels of qualitative variables (CTRL, DMSO and PCA) and hence, multiple-way ANOVA or Kruskal Wallis tests were performed when the hypothesis satisfied conditions of normality or not, respectively. The significance level (α) was set at 0.05 and significant values were represented in the graphs as: ^#^ 0.09>*p*>0.05; \**p*<0.05; \*\**p*<0.01; \*\*\**p*<0.001.

The ICA is a higher order statistics and the number of FP generators returned may vary from one sample to another depending on the animal’s processing demands. We rejected all components that contributed less than 1% of the total variance.^37^ The mean power of the component or their frequency content was compared for each of the treatments (Control 3 min, DMSO 30 min, PCA 40min) and among individuals (n = 6 animals). The same statistical tests were used as indicated above.

## Results

### Systemic delivery of protocatechuic acid enhances Schaffer-CA1 excitatory transmission

Our initial studies set out to explore whether systemic delivery of PCA had any effect on electrophysiological parameters in the hippocampus of anesthetized animals, assessing evoked potentials in a well-characterized excitatory pathway, the Schaffer input to the CA1 region (Figure 2(a)). We chose an intensity that elicited an o-PS about one third of the maximum, which allowed amplitude changes in any direction to be detected, although it also yielded significant spontaneous fluctuations. All the responses were averaged over one minute periods (n = 6). After a 3 min control recording, DMSO (1%) was injected (i.p.) and 30 min later, a single dose of PCA (150 mg/kg i.p.) was delivered. While DMSO produced no significant effect on the fEPSP or the o-PS, PCA significantly increased both potentials (expressed as the % of the control 3 min post injection): fEPSP, 114.6 ± 17.1 DMSO vs. 128 ± 30.2 PCA (n = 12, *p* = 0.347, Mann Whitney); o-PS, 115.4 ± 19.2 DMSO *vs*. 128.1 ± 31.2 PCA (n = 12, *p* = 0.219: Figures 2(b),(c)). A mild rundown was associated with this effect that reduced the statistical significance after a few minutes, although the enhancement was evident up to 30 min post injection.

**Figure 2.**
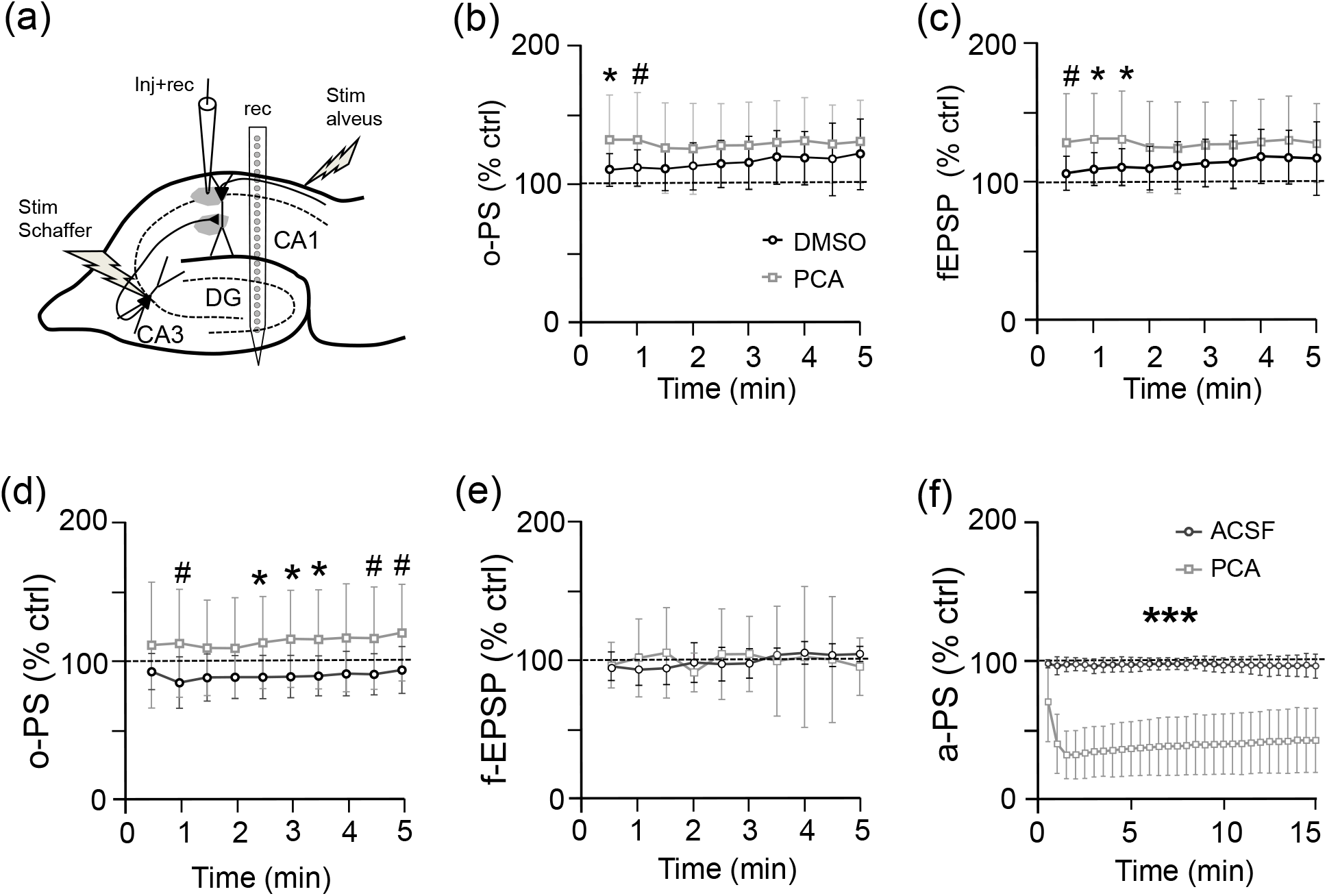
Scheme of the electrode arrangement (a), and of the systemic (b,c) and direct (d-f) effects of PCA on CA1 evoked potentials. To explore the systemic effects of PCA, a linear array (rec) was used spanning the lower cortex, CA1 and DG. Stimuli were delivered to the CA3b for Schaffer activation (stim Schaffer) or the alveus (stim alveus) for antidromic activation. To test the acute effects of PCA, a drug-filled pipette was used that simultaneously recorded and performed microejection (Inj+rec), either in the cell body layer or the st. radiatum (shaded areas). (b) The effects of a single intraperitoneal PCA injection on the o-PS (b) and f-EPSP (c) at optimal sites of the linear probe. The two responses were obtained from separate trials, adjusting the intensity to obtain 30% of maximal response in each case. The time points represent the mean ± s.e.m. of n= 6 responses and as a control, the vehicle (DMSO) was injected in another group of animals. A mild increase of the two potentials was observed after PCA administration that reached a significant level in the first minutes. (d), (e) and (f) plot the direct effects of ACSF or PCA microdrops on the a-PS, o-PS and fEPSP at the respective loci. The o-PS but not the f-EPSP increased slightly, while the a-PS diminished dramatically and in a sustained manner. All the data are normalized to the corresponding pre-injection values: CTRL (n=5×2), ACSF (n=5×2), PCA (n=5×2).

### Paradoxical effect of intraparenchymal protocatechuic acid on the ortho- and anti-dromic somatic population spike

The initial findings were compatible with an enhanced o-PS in CA1 following the synaptic response. However, since PCA crosses the BBB,^15^ it can affect hippocampal excitability in multiple manners and through different pathways. Hence, we assessed if PCA had any direct effects on domain-specific electrogenic events of the same neuron population by performing pressure microejection from the recording pipette (Figure 2(a)) onto different cell domains clustered in the CA1 strata.

PCA microinjection into the st. pyramidale did not significantly affect the o-PS relative to the pre-injection values (expressed as the % of the control: 99.8 ± 12.7 *vs*. 116 ± 35.3, n = 9, *p* = 1 Mann Whitney) (Figure 2(d)). Similarly, fEPSPs did not vary significantly when PCA was delivered to the st. radiatum (Figure 2(e)). At equivalent times post injection, microdrops of the vehicle (ACSF-DMSO) tended to reduce the o-PS (88.5 ± 15.4), which was significantly different than the values obtained after PCA administration (*p* = 0.021 at 3 min post-injection, Mann Whitney). We also explored the effect of PCA on spike electrogenesis following antidromic stimulation of the alvear tract (Figure 2(f)). The a-PS was not affected by vehicle injection into the st. pyramidale (101.3 ± 2.1 vs 95.8 ± 5.4, before and 3 min after injection: n = 10, *p* = 0.28 Mann Whitney), yet PCA caused a rapid and strong decline in the a-PS (99.4 ± 5.7 *vs* 34.1 ± 18.2: n = 10, *p* = 0.0001 Mann Whitney). This effect persisted and it had barely recovered 40 min post-injection (42.2 ± 23.2 at 40 min post-PCA as opposed to 97.4 ± 4.0 15 min post ACSF: n = 10 in both cases). The lack of an effect on the o-PS indicates that neurons remain electrically viable after PCA administration and that they may still produce synaptically elicited feed-forward spikes (dendrite-to-soma).^29,41^ By contrast, the strong reduction of the a-PS suggests a dysfunction of the spike-generating zone at the axon-soma junction.

### Systemic delivery of protocatechuic acid produces a lasting and varied change to the spontaneous activity at specific FP generators

In light of the above, we explored if systemic delivery of PCA might produce persistent alterations to ongoing activity in the hippocampus and cortex by examining pathway-specific FP generators, these established from multisite spatial maps of FPs using the ICA. FP generators avoid the mutual distortion of waveforms inherent to raw FPs and hence, drug-induced changes can be more accurately explored over time.^25^ The site-dependency of raw (multisource) FPs recorded from the cortex (upper traces) down to the DG was analyzed, extracting FP generators through the ICA (Figure 3, colored traces and profiles). There were differences between the waveforms and patterns of raw FPs, and those of the isolated FP generators, which were more prominent at some sites rather than others, reflecting the power at different sites (see spatial voltage profiles at the top right panel). As such, we examined the following main FP generators identified through their characteristic profiles^34,38^: the Schaffer input to CA1 (Sch); the input to the st. lacunosum-moleculare (L-M) in the CA1; the Perforant Path input from the entorhinal cortex to the DG (PP); an inhibitory input to granule cell somata (GCsom); and the main FP component in the cortex (Ctx).

**Figure 3.**
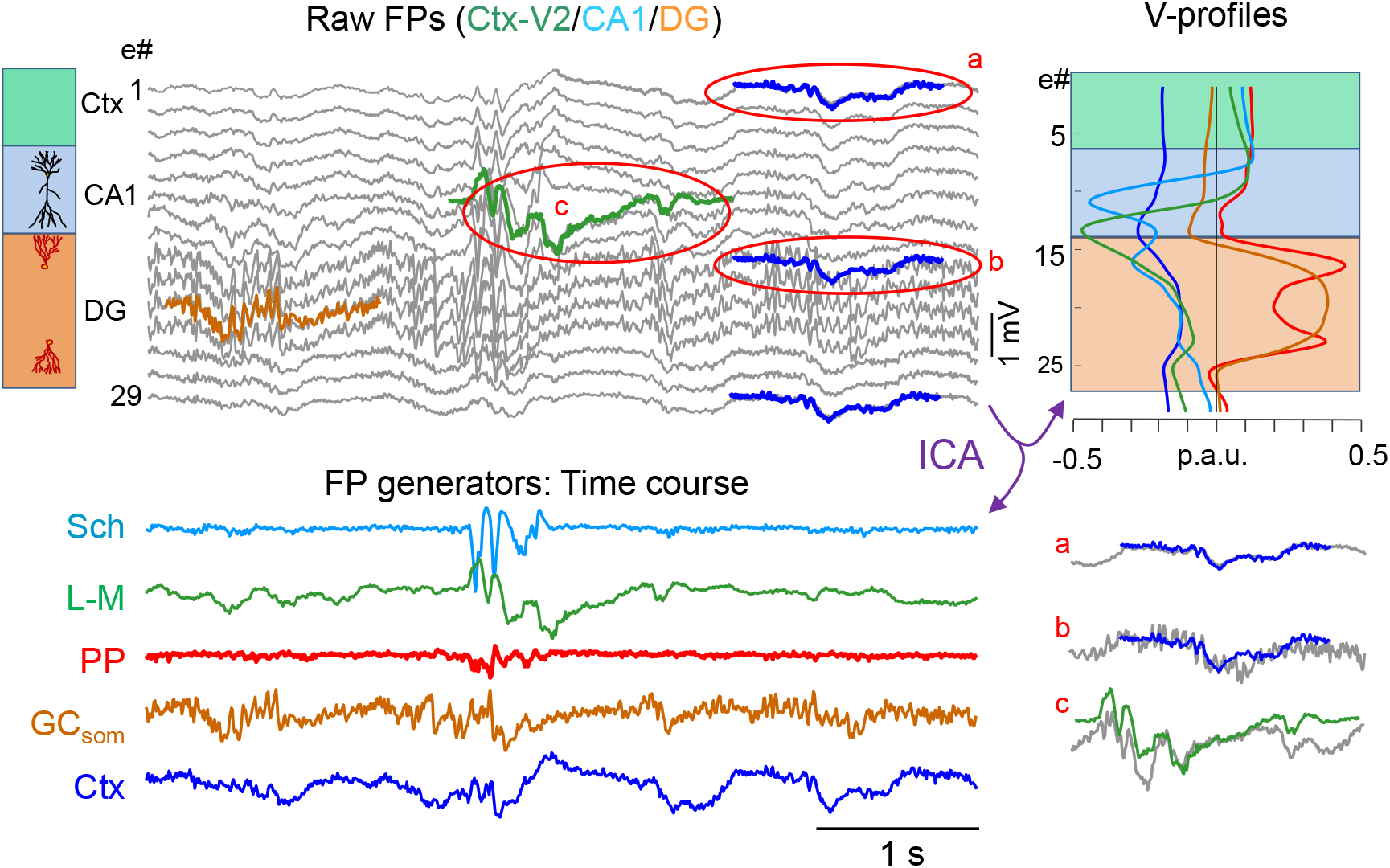
Disentangling the FP generators to access the pathway-specific time courses. Sample epochs of raw FPs (upper traces in grey) recorded using a linear array across the V2 cortex and the CA1/DG (only every other channel is displayed), and showing the site-dependence and multifarious collection of FP waveforms. The ICA returns spatially-coherent components (FP generators) and provides readout of the temporal dynamics free from the contribution of others (color-coded traces). The five main FP generators separated are labelled according to their maximal anatomical layers: Sch (CA1 Schaffer), L-M (CA1 st. lacunosum-moleculare), PP (perforant path input to DG), GCsom (somatic input to granule cells), and Ctx (main cortical generator). The amplitudes are presented in proportional arbitrary units (p.a.u.) and the corresponding spatial distribution (i.e. the relative power along the recording track) is shown on the right (voltage [V]-profiles). A few instances are superimposed onto the raw FPs to facilitate their visual matching/unmatching. The better the FP generator matches the raw FP at a particular recording site, the larger its relative contribution, as illustrated by the parts of the recordings highlighted by red ovals and extracted below: (a) a slow wave in the Ctx generator (dark blue) tightly matches the raw FPs in upper (cortical) channels (in grey), but it also spreads by volume conduction all the way down to the DG (b), where it accounts for the slow envelope of the FP there, although at this site there are also faster superimposed oscillations generated at other sites. This justifies the use of FP generators as opposed to raw FPs to quantify electrical activity: e#, electrode number: only every other channel is displayed.

The dynamics of the basal electrical activity were quite variable, similar to wakefulness but with dominant patterns characteristic of anesthesia. For example, there is slow wave activity in the cortex and irregular activity interspersed with epochs of gamma, alpha or beta oscillations in specific sub-fields and strata of the hippocampus.^42^ Over an elongated time scale, FP activity appeared as intermittent bursts of strong and weak power that last several minutes (Figure 3(a)). The pattern and duration of these periods was heterogeneous between individuals, although the mean power was generator-specific and remained within fairly stable limits in each animal. The latter condition was considered the minimum stability criterion for validating drug effects. The existence of different FP patterns in control conditions may raise doubts as to whether sufficient standardization of the data can be achieved and indeed, they pose difficulties in choosing quantifiers of the drug effect. This became necessary as the onset of the effects turned out to be gradual, long lasting but varying in strength across the population. Therefore, we used different quantification strategies to account for the multiple time-scales in which FP activity develops, i.e. individual waves, patterns and trends.

Systemic i.p. injection of PCA produced changes in three of the five animals, evident through compacted temporal displays of the power at which local peaks are maintained (Δt = 5 s: Figure 4(a)). This means that at least some waves were larger than average after PCA injection. Alterations appeared 2-5 min after injection, they peaked after 10-15 min and then faded over 30 min. In comparison to ACSF or DMSO injections, the PCA-induced changes in combinations of the following: the regularity, duration, amplitude, temporal structure, or baseline level of activity. Notably, these affected some FP generators but not all, and not in all animals.

**Figure 4.**
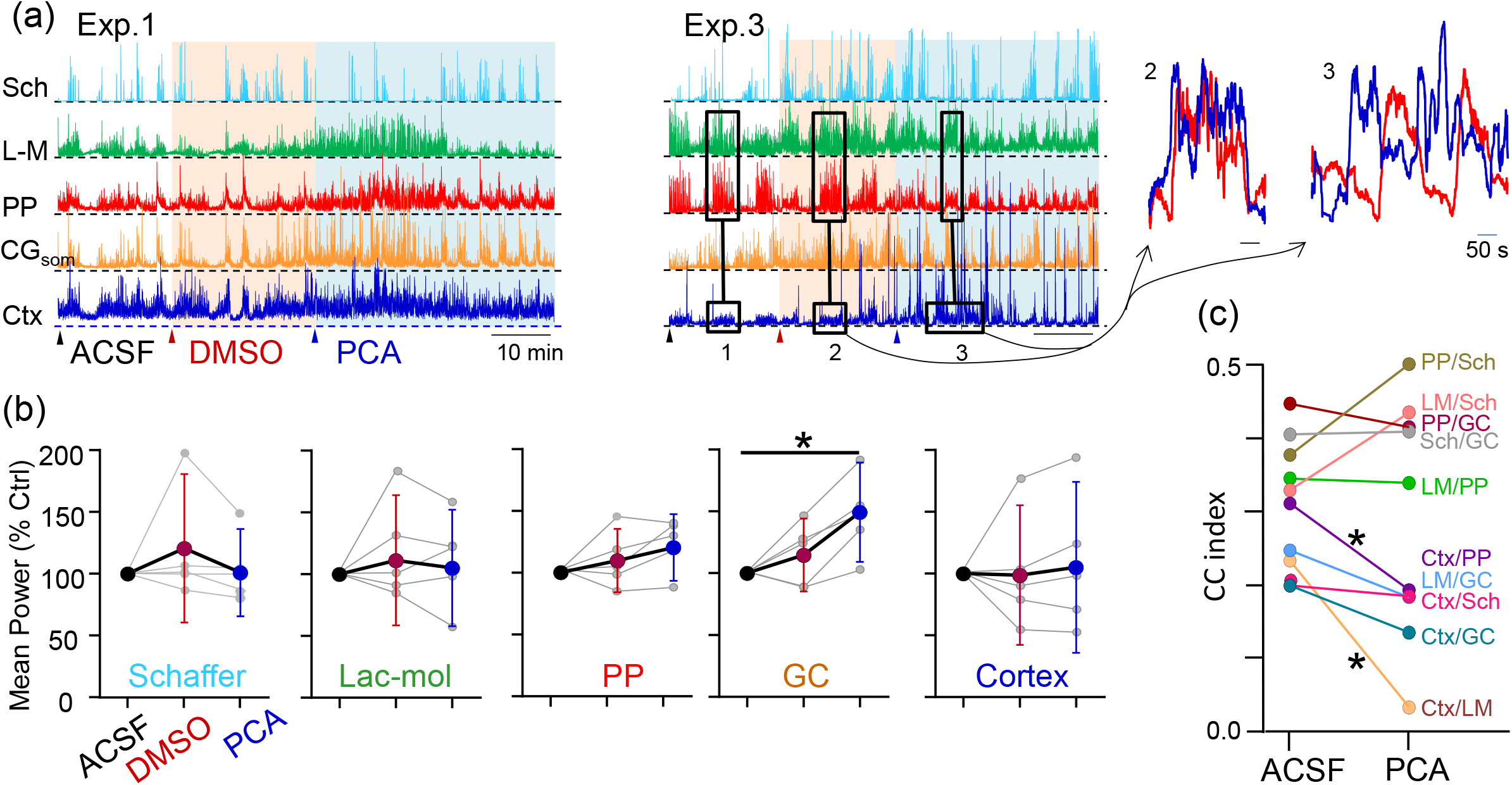
PCA has a variable effect on the FP generators and it produces cortico-hippocampal disengagement. (a) Time plots of the power of the separate FP generators in two experiments (arbitrary units). Compact plots maintain the peak values in which the colored backgrounds mark the periods after each injection (arrowheads). Although the resting patterns of activity (ACSF and DMSO) could differ amongst animals, there is clearly tight covariance between the cortical and hippocampal generators (except the Schaffer). Note the graded and selective action of PCA on some generators and the disruption of cortico-hippocampal covariance. The epochs marked by the black boxes 2 and 3 are amplified on the right for two generators. (b) Mean power and standard error for each FP generator evaluated over epochs of 15 mins after each injection. The values from individual animals are in grey and the data are normalized with respect to the ACSF. The strong variability amongst animals hindered the statistical analysis. (c) The cross-correlation (CC) index between pairs of generators grouped across the population (n = 5) highlighted a drop in the CCs between the cortical and hippocampal generators (except the Schaffer).

We first used descriptive quantifiers to characterize spontaneous activity, such as the mean power (Figure 4(b)). When this was averaged over 15 min periods, we found a wide spread among animals, even when the data were normalized relative to the control (ACSF). Such dispersion resulted in a poor population effect of the drug, as witnessed by only one FP generator reaching statistical significance (GCsom), although clear changes in other generators were observed in some individuals. We then examined the time courses and power envelopes at higher temporal resolution, and we found that the peak values in the mean power plots belonged to larger than normal waves or wave groups, yet it did not appear that their frequency had a strong impact on the mean power.

An interesting observation was that an alternation of high and low power episodes was strongly correlated between the cortical and most hippocampal FP generators in the control situation (ACSF or DMSO administration: Figure 4(a) Exp#3, boxes 1 and 2). However, delivery of PCA disrupted this temporal relationship between the generators of different structures (box 3), although it remained strong internally among the hippocampal sub-fields and generators. The cross-correlation (CC) index between power envelopes for pairs of generators confirmed this (Figure 4(c)). For instance, the mean Ctx/L-M CC was 0.23 ± 0.06 after ACSF administration but it dropped to 0.03 ± 0.05 after PCA delivery (t = 6.56, *p* = 0.001; Student t-Test, n=5 animals), whereas the L-M/PP CC was 0.35 ± 0.11 after ACSF (ctrl) and 0.34 ± 0.14 after PCA (t = 0.07, *p* = 0.47). Hence, PCA appears to produce some type of functional detachment of the cortex and hippocampus.

We then tested the effect of PCA on the frequency content of specific FP generators, which highlighted significant differences between the average power after ACSF and PCA delivery in 10 of the 25 possible comparisons (5 generators and 5 frequency bands: Supplementary Figure 1). These were mainly clustered in the L-M, GCsom and PP generators of the hippocampus, and within medium-high frequency bands (theta and beta), although some generators were also altered in the alpha and gamma bands. In all of these, the mean power increased after PCA delivery. Notably, an increase in the mean power was also evident after DMSO administration in four comparisons. The cortical FP generator was barely altered in any frequency band and it also presented the weakest variability. A dominant contribution of low frequencies to the overall power in the wideband comparisons (Figure 4(b)) may explain the failure of these to reveal population effects.

Since some of the alterations produced by PCA injection had little impact on the mean power over longer periods, such as the changes in the amplitude of individual waves, we devised two additional methods to optimize their detection. First, we estimated the number of waves above a series of voltage thresholds (with standard deviations: 0.5xσ, σ, 2xσ and 3xσ). An illustrative example of the waves detected at different voltage thresholds and following distinct treatments can be seen for the Sch and L-M. generators in a representative animal (Figures 5(a) and 5(b), respectively). In individual animals, PCA transiently modified the number of waves per second in some FP generators (epochs shaded in blue), an effect that was unmasked as the threshold for detection increased, although these observations were not clearly appreciated in the population analysis. Thus, the cortical generator in the high frequency band reached a significant level for all but one of the possible comparisons (*p* = 0.002). We then compared the effect of PCA injection on the number of waves per generator and voltage threshold (Ctrl *vs* PCA), which turned out to be not very discriminating as only one generator returned a significance difference (PP generator at σ = 1, *p* = 0.016, U = 24; Mann-Whitney test).

**Figure 5.**
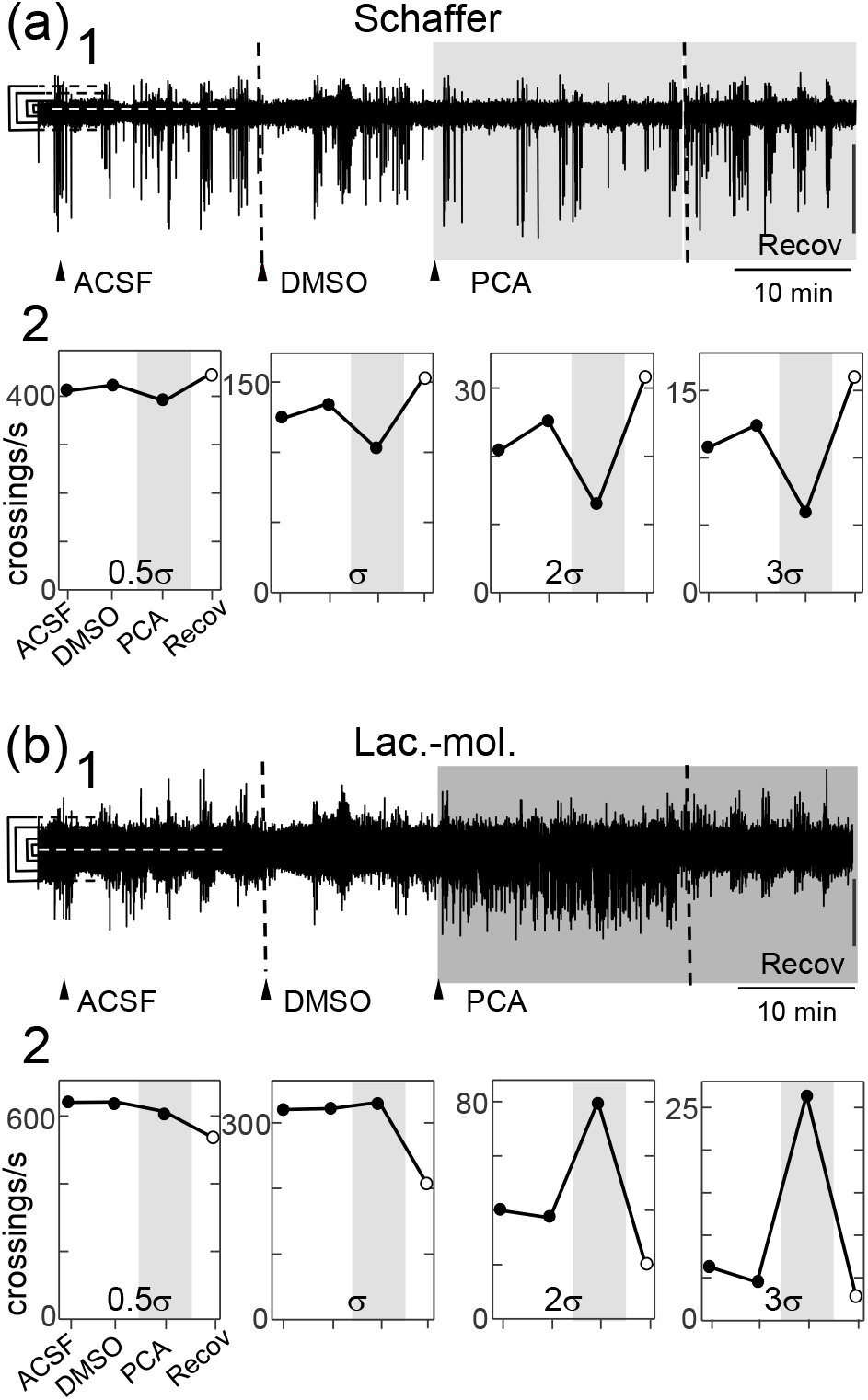
The effects of PCA on the number of waves in FP generators at different voltage thresholds. The mean value and the thresholds are indicated to the left, 0.5 (green), 1 (cyan), 2 (red) and 3 (black) times the value of the standard deviation of the mean. (a), (b) correspond to the Schaffer and Lac-Mol generators in the same animal, respectively. The upper traces (a1, b1) show the time course of the activity over a compact scale. The number of waves per second is plotted below (a2, b2) for each voltage threshold. The voltage in the FP generators was estimated at the site that presented the maxima. At the Schaffer generator, the bulk of the waves belong to small amplitude (50-200 μV) gamma waves contained in the thick baseline, whereas the large strokes belong to sharp-wave complexes. In the L-M generator, the duration of the waves had a greater dispersion, giving a more irregular global pattern. The longer waves induced by PCA were mostly delta-type slow waves.

Different proportions of long and short duration waves in the control and PCA epochs may bias the analysis, since fast small-amplitude waves may not be detected when riding on slow ones. To minimize this possible bias, we re-analyzed the data after splitting the time course of the FP generator into two frequency bands above and below 10 Hz. Thus, one contained mostly delta and theta activity, while the other was predominantly in the beta, alfa and gamma bands. This procedure optimized the counting of the waves, and some differences in the effect of PCA were readily appreciated in certain individuals (Supplementary Figure 2). However, the population statistics rendered similar results as the wideband analysis.

A second approach was to compare the density distributions of the time-values for each FP generator after ACSF and PCA delivery (grey and cyan areas in Figure 6). We estimated the mean, standard deviation, kurtosis and skewness of the distributions, and to check the stability of these parameters over time with each treatment, they were segmented in 90 s consecutive epochs and normality was tested for each parameter (47 out 100 comparisons: 5 animals x 5 generators x 4 parameters). The results obtained were heterogeneous and again indicated differences between some generators in some animals (Supplementary Table 1). Globally, 55 comparisons reached a significant level, of which only 30 presented a normal distribution. The parameters of the density distribution that reached significance in the ACSF-PCA comparisons were (in decreasing order), the mean (n = 19), standard deviation (n = 16), skewness (n = 12) and kurtosis (8 of a total of 25 in each case). The generators that achieved significance were (# of significant comparisons in decreasing order), the Cortex (n = 14), L-M (n = 13), GCsom (n = 12), Sch (n = 8) and PP (n = 8, of a total of 20 each). All five animals presented PCA induced differences relative to control conditions (# of significant comparisons for the five animals: 14, 9, 11, 8, 13, out of 20 possible each).

**Figure 6.**
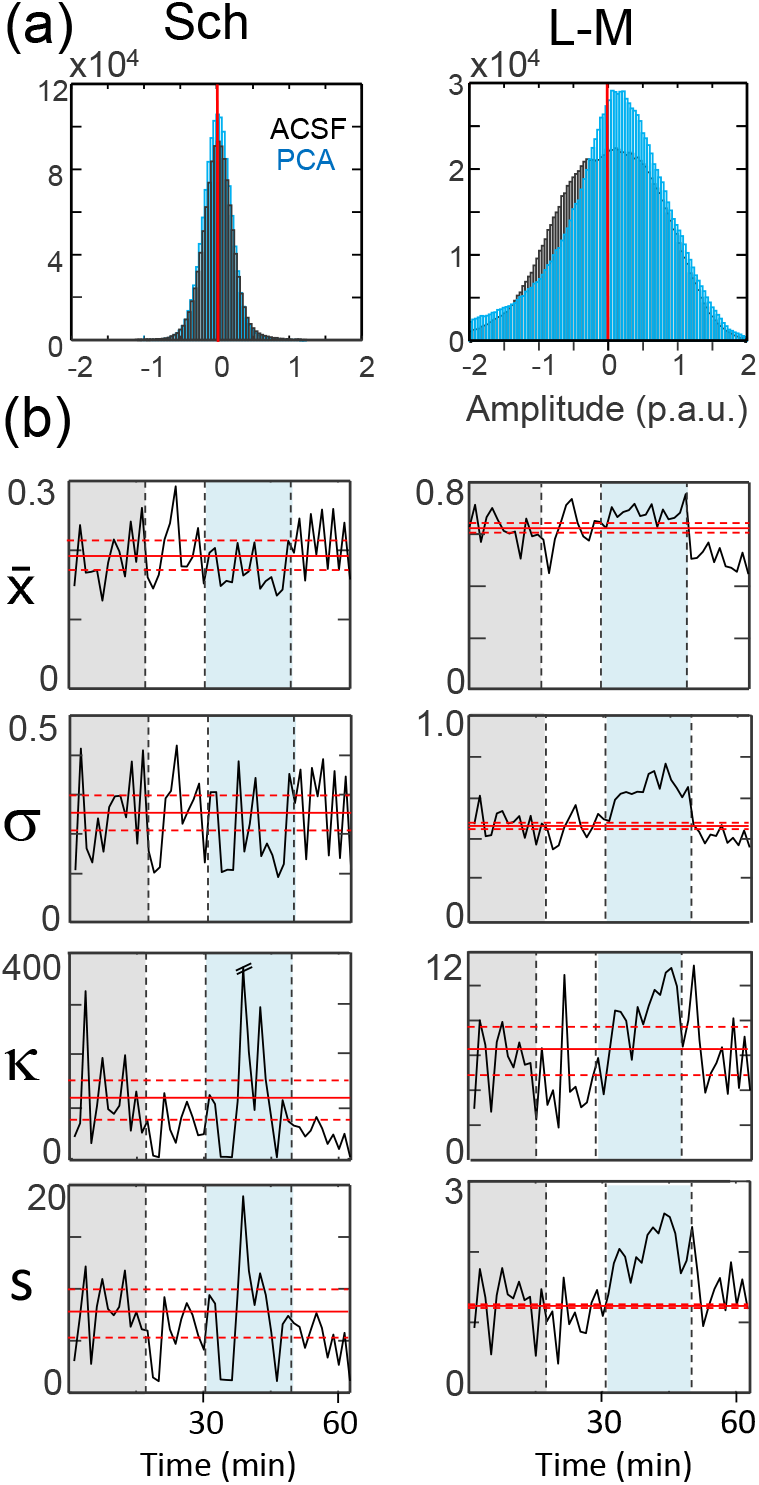
Density distributions of the time-point voltages efficiently unmasks the effects of PCA on the time-course of FP generators. The parameters analyzed were the mean (), standard deviation (σ), kurtosis (κ) and the skewness (s). The left and right columns show a representative example for the Schaffer and Lac-mol generators in one animal. (a) Superimposed distributions of time point voltages after control (ACSF) and PCA (blue) delivery. (b) The epochs were split into consecutive 90 s fragments and the distribution parameters plotted as successive values (black lines). The mean and 95% confident limits were estimated for the control period (grey shadow) and are represented by horizontal red lines expanded to all treatments to facilitate the visual comparison. In the example illustrated, the values characterizing the distribution for the Schaffer generator were about the same in control and PCA (blue shadow), whereas PCA modified the standard deviation and the skewness for the Lac-mol generator.

### Systemic protocatechuic acid has no effect on memory-related hippocampal-dependent behavioral tests

To explore whether the changes in electrical activity induced by PCA translates into a behavioral manifestation, we checked the performance of rats in two different behavioral tests (Supplementary Figure 3). First, when the animals were subjected to the NOR test, those that received saline + DMSO or PCA i.p. learned efficiently in the training period and both performed similarly during the test. There was a tendency towards worse performance in the PCA-treated group, although this did not reach the accepted level of significance (% discrimination 47.55 ± 11.65 *vs* 33.4 ± 13.65, t-test independent samples, *p* = 0.116, n = 5). We found similar results for the object location test (OL: % of discrimination 40.73 ± 16.26 *vs* 36.88 ± 20.85, t-test independent samples, *p* = 0.753, n = 5), such that both groups were able to discriminate which of the two objects moved, yet there were no differences between the animals administered PCA and those that received saline.

## Discussion

The present study shows that the phenolic compound PCA alters electrophysiological activity in the hippocampus and cortex of anesthetized animals, and that these effects vary depending on the route of administration. We report acute and sustained effects of PCA, particularly in the ongoing activity of pathway-specific FP generators. Notably, we found a detachment of overall cortico-hippocampal coherence. Accordingly, we propose that some polyphenols exert a direct neuromodulatory-like action on central brain structures.

### Technical considerations

The FP analysis presented here highlights a number of caveats derived from the multisource and spatial nature of these signals,^32,43^ even though many were minimized by using an ICA that extracts spatially coherent components or FP generators.^33,44^ This step is important to provide more realistic time courses by avoiding mutual contamination between spatially overlapped sources or with potentials originated in distant sources (volume-conduction), both of which affect measurements like the mean power or the frequency content. The spatial gradients and landmarks of the generators obtained in this study were those reported previously, which helped their identification. However, the diverse activity patterns common to each FP generator^34,36^ posed a challenge for their quantification and normalization between animals. This could have been optimized by using a deeper anesthetic plane to achieve a more uniform electrographic pattern.^45^ However, we chose to collect data as close as possible to awaken animals so as not to limit the potential utility of the study. Among the quantifiers used, the mean power or number of waves of different amplitude proved to be particularly sensitive to the high variance of the population data, whereas splitting the FP generators into frequency bands or using density distributions of time-point voltage was more effective for the present data set. A case-by-case discussion of the suitability of each approach is beyond the scope of this study, yet some facets are evident. For example, there was clearly a significant change in the number of large amplitude waves at some FP generators, although the relatively low number of these waves means they are unlikely to alter the mean power over long epochs.

### The cellular actions of PCA appear to be non-specific

The short duration of the increase in fEPSPs along the CA3-CA1 pathway upon direct PCA delivery makes it unlikely that this is related to synaptic plasticity and moreover, systemic delivery of PCA leaves fEPSPs unchanged. Without additional information, this apparent paradox may have many explanations, such as differences in the concentration of PCA that reaches the target site depending on the route of administration, or the multiplicity of actions at different cellular targets that become balanced at any particular synapse studied. Similar neuromodulatory-like actions of orthodromic responses have been observed for various neuroactive compounds, such as activators of the type 1 CRH receptor and P2 purinergic receptors, or some protein kinase C inhibitors.^46-48^ As such, future studies should address whether the effects of PCA on evoked activity interact with these or other receptors/signaling pathways. It is important to note that a small or even null effect on evoked potentials does not rule out changes in the ongoing activity in the pathway stimulated (see below).

Other findings in this study are compatible with non-specific membrane actions of PCA, such as the persistent and strong reduction of the a-PS upon direct delivery to the soma layer. Injected volumes might experience depression as some channels and receptors display strong mechanosensitivity,^49^ yet we ruled out this possibility by studying volumes previously found not to alter evoked activity.^50^ Larger drops produce transient depression on somatic a-PSs (1-5 min), whereas here this depression persisted for 30 minutes. Moreover, ACSF or DMSO injections had little or no effect. A direct anesthetic-like effect of PCA is also unlikely, such as that produced by lidocaine that is evident through a blockade of Na^+^ channels,^51^ since the orthodromic PS remains unchanged or enhanced. One possible explanation for this is that the PCA destabilizes membranes at sites of high Na^+^ channel density, such as the axon hillock.^52^ This highly specialized membrane may impede axonal spikes from invading the axon hillock soma junction for long periods,^53^ for instance if anchoring protein re-assembly is required. Such a possibility is compatible with the maintenance of the o-PS, which initiates in dendritic loci in vivo and propagates forward to the soma.^29^ Very few compounds other than Na^+^ channel blockers affect a-PS but there are some, like serotonin, that reduce a-PS trains through an increase in conductance.^54^ Further exploration using *in vitro* preparations should help clarify the direct effects of PCA and whether there is neuron-selectivity. In particular, it will be important to elucidate whether these effects are mediated by a receptor or if they are membrane unspecific. Some reported effects of PCA, such as its antioxidant properties,^55^ may play such a role altering the physicochemical properties of the membrane or the extracellular matrix. It will also be important to explore the range of effective concentrations of PCA in order to establish appropriate ranges for any beneficial/detrimental effects of this compound.^14^

### Cortico-hippocampal disengagement by PCA

One important finding of this study that is robust in terms of population statistics is the PCA-induced drop in the CC between the cortical and hippocampal FP generators, while the CC between hippocampal subpopulations is not affected. In principle, this observation suggests a functional disengagement between the cortex and hippocampus. Since FPs are convolution of synaptic activity it might be argued that these hippocampal generators also reflect cortical (entorhinal) input, in which case such disengagement would be better defined as cortico-cortical, i.e. among different cortical areas. Indeed, it is known that slow-wave activity in the cortex may spread internally or that some regions may be sidelined.^56^ A reduction of the overall cortico-hippocampal CC index may arise after changes to the dominant electrographic pattern, shifting from that based on delta activity to another based on fast activity. One such change characterizes the transition from sleep to awakening.^57^ A similar reaction can also be achieved through enhanced sensitivity to brainstem inputs, which are known to be modulated by polyphenols.^58,59^ In some individuals we saw an intermittent transition from slow-wave activity in the control towards a desynchronized state after PCA delivery, although this could not be generalized due to the variation in the resting patterns among the population.

### PCA fails to modify hippocampal-dependent tasks

In our hands, the PCA had little effect on two hippocampal dependent behavioral tests, which contrasts with observations in the literature reporting improved performance in tests assessing memory performance.^60-63^ Differences in the protocols may underlie the contrasting results. Thus, former studies explore memory alterations under pathological conditions (hypoxia, Alzheimer’s, etc.), which may potentially reflect protection provided against memory loss rather than PCA producing true improvement. Furthermore, treatment was delivered over several days in most studies, implying the administration of a higher dose and different patterns of PCA administration to that delivered here. Notably, a study in healthy adult rats exploring the effects of repeated PCA administration over several days on hippocampal-dependent memory rendered similar results to ours.^64^ Yet, memory formation is a very complex process involving numerous brain regions,^65-67^ hence this issue clearly requires further study.

### A hypothesis based on the time scales and infra-slow oscillations

In the present study we addressed the acute electrophysiological effects of PCA administered in a single-dose protocol, observing effects lasting from a few to tens of minutes. However, the literature available reports the long-term benefits of PCA due to possible antioxidant, anti-inflammatory,^68^ immunoregulatory^69^ and even neuroprotective effects.^13^ In this context, our results should be taken with caution, and the effects of PCA on additional cerebral and systemic targets should be explored over different time scales.

In relation to this, the high level of coherence between hippocampal and cortical generators under control conditions occurs over a very long time scale. The mechanisms underlying this global coherence between structures probably extend beyond the phenomena of pacemakers and clocking that set standard network oscillations. Infraslow brain activity is reminiscent of the resting state fluctuations reported in humans,^70-72^ although the underlying mechanisms are as yet poorly understood. In animal models, some ultraslow potentials are related to glial-mediated network synchronization.^73^ Indeed, it would be interesting to check the effects of polyphenols on glial cells, since PCA is known to produce multiple neuroprotective effects in some pathologies. For instance, PCA decreases the levels of oxidative stress^74,75^ and neuroinflammation,^76,77^ events in which glia are known to participate.^78^

### Concluding remarks

This study shows that PCA directly targets the hippocampus, and that it has both acute and sustained actions on cell and network electrophysiology, as well as on global coherence between structures. Some of the results point to non-specific effects and thus, we suggest futures studies should also investigate the actions of polyphenols on cells other than neurons. Non-specific non-neuronal effects make it likely that polyphenols could interact synergistically with individual or environmental conditions, such as the animal’s metabolic state or genetic factors. We propose to take the present results as evidence for the direct effects of a dietary polyphenol on the electrical activity of neurons, with the hope that future studies will specifically address how these compounds influence physiology and pathologies over multiple time scales. The results presented here also indicate the importance of the route employed for polyphenol delivery, which should be taken in account in order to better identify which effects are more likely to be produced following natural dietary intake.

## Acknowledgements

We thank Mark Sefton at BiomedRed for editorial assistance. OH and JM belong to the Cajal Blue Brain Interdisciplinary Platform of the CSIC, Spain.

## Author contribution statement

MMA performed experiments, MMA, RMA, and JM, analyzed data, BB, MVMA and OH designed experiments and edited the manuscript, OH wrote the manuscript.

## Supplemental material

**Supplementary Figure 1.**
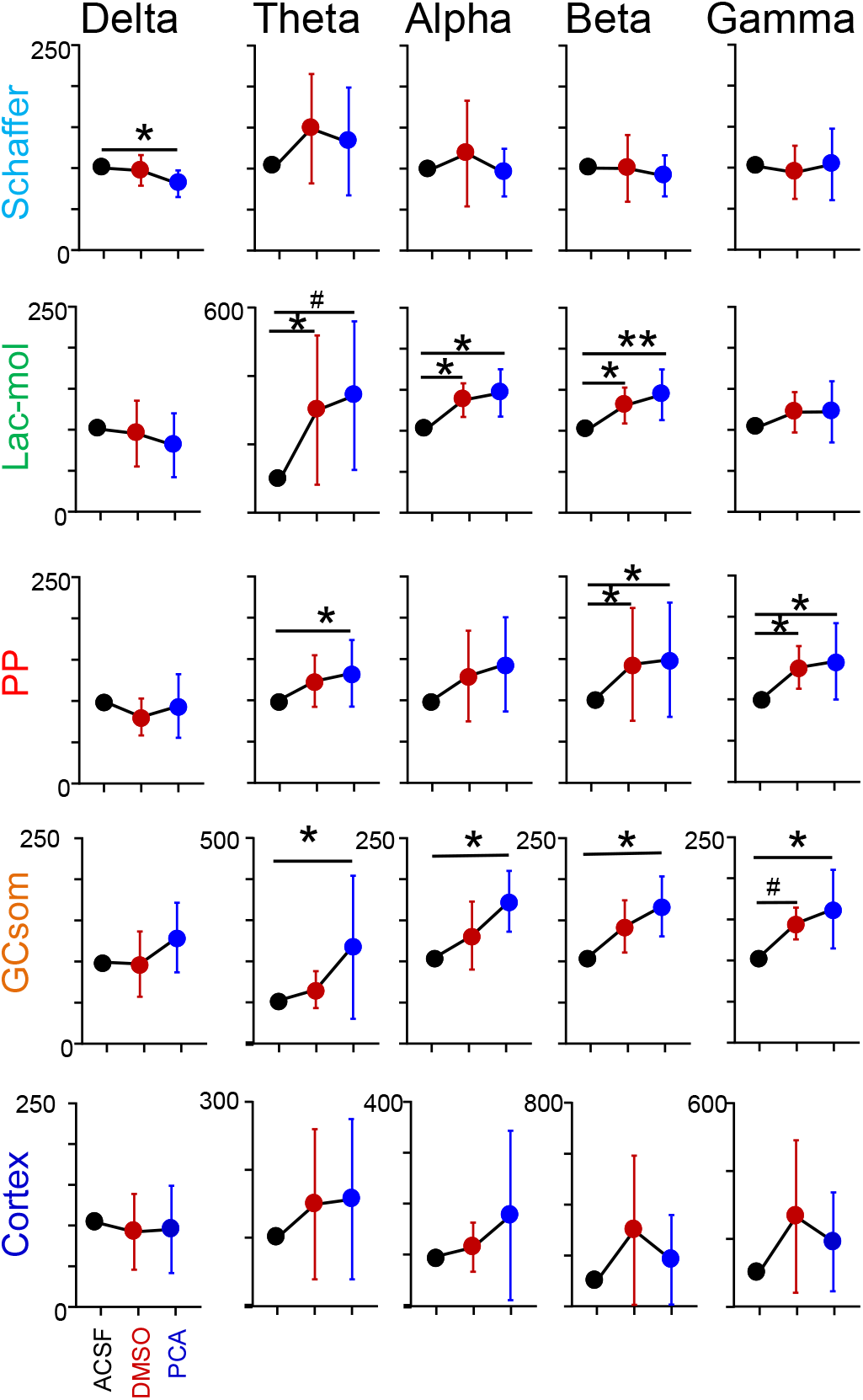
The effect of PCA on the frequency content of the different FP generators. The values represent the mean power and standard error (n = 5) normalized to the control (ACSF, black dots) averaged in 15 min epochs after each treatment. PCA produced a significant increase in the population values for 10 of the 25 comparisons (Student t-test). Significant changes were concentrated in the L-M, PP and GCsom generators, and in the mid-high frequency bands (theta, alpha, beta). Therefore, focusing on the frequency content of each FP generator was more effective in discriminating the effects of PCA than when estimated over wideband power.

**Supplementary Figure 2.**
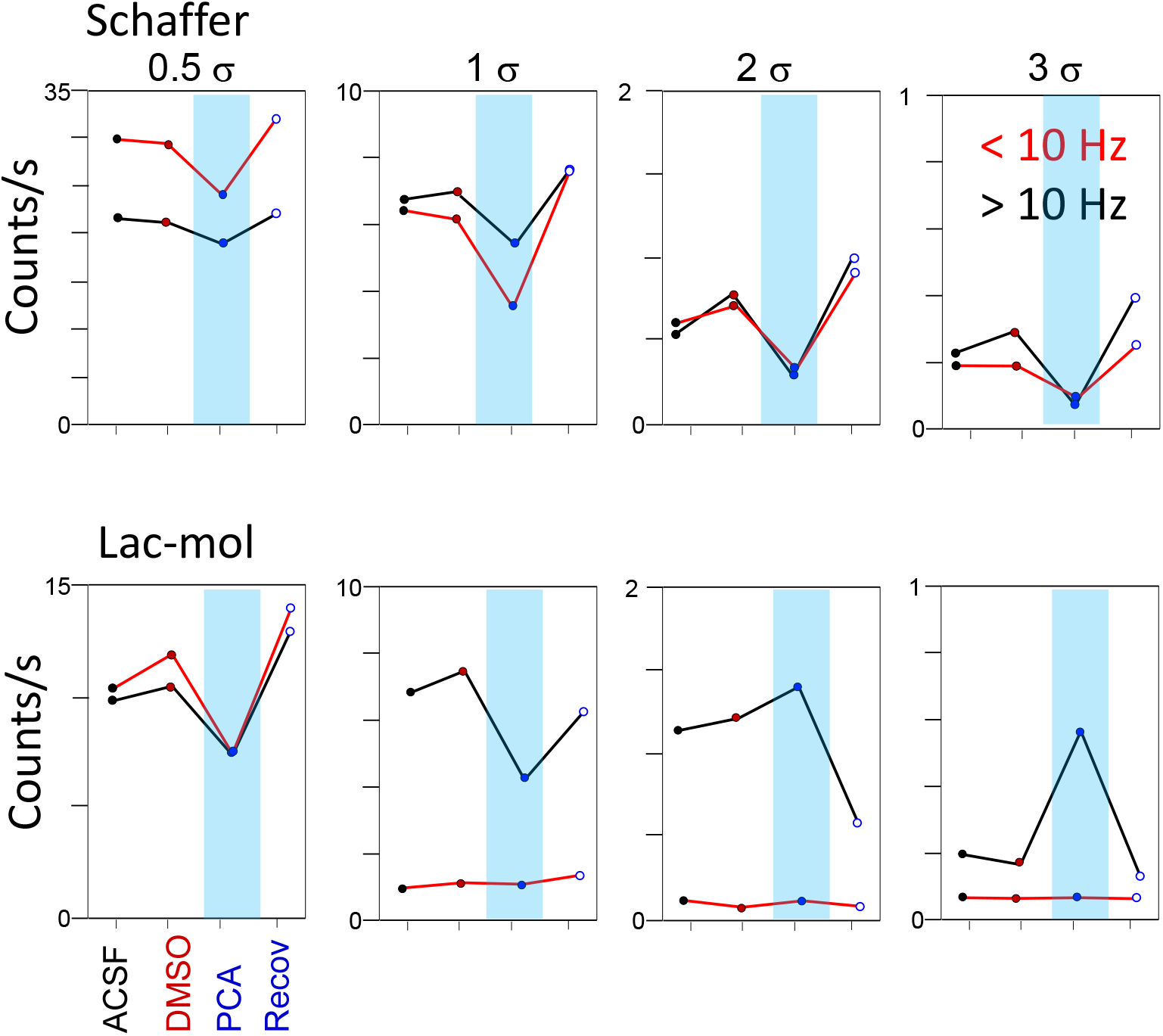
Re-analysis of the data for the Schaffer and the L-M generators of a representative animal (see Figure 5). The time course of the FP generators was filter-split into two sets of high and low frequency bands (10 Hz cut-off frequency) to avoid omitting small fast waves that ride on top of slower ones. This effect can be appreciated in the small number of high frequency waves in the L-M (fast waves are generally smaller). PCA had similar effects as on the wideband signal, yet some differences were seen. Note PCA reduced the waves detected in the lower voltage thresholds (0.5 σ and 1σ) but it increased those in the wideband analysis of Figure 5 at any threshold.

**Supplementary Figure 3.**
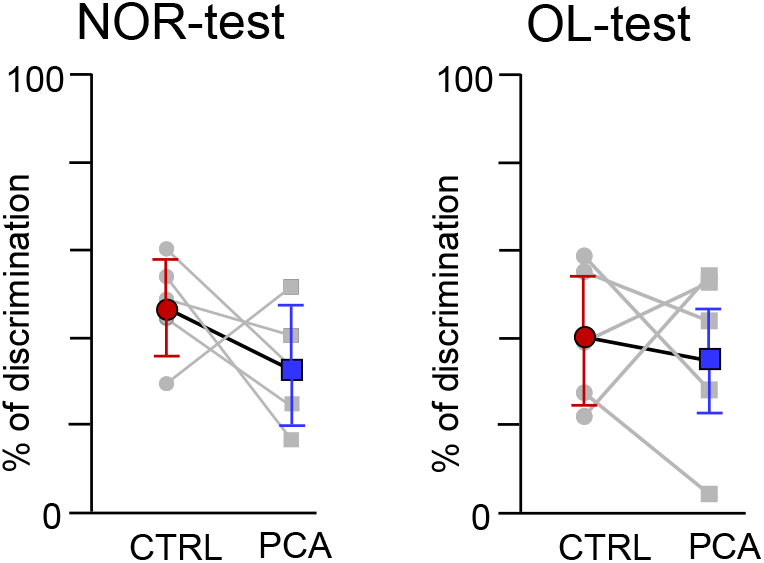
PCA has little impact on hippocampal-dependent tests based on the results obtained for individual animals (in gray), and the mean and standard error in color: NOR, novel object recognition; OL, object location. All but one animal showed reduced performance in the NOR test and the mean did not reach a significant level.

**Supplementary Table 1.** Statistical analysis comparing the parameters of the density distributions of the voltages per rat and FP generator after ACSF and PCA delivery. For normal distributions we used a Student t-test and a non-parametric U-Mann Whitney test for the others. Significant *p* values are marked in red and the alpha is 0.05 in all cases: 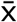, mean; σ, standard deviation; κ, kurtosis; Sk, skewness.

|  |  | Rat #1 |  |  | Rat #2 |  |  | Rat #3 |  |  | Rat #4 |  |  | Rat #5 |  |  |
| --- | --- | --- | --- | --- | --- | --- | --- | --- | --- | --- | --- | --- | --- | --- | --- | --- |
|  |  | *t* | *U* | *p* | *t* | *U* | *p* | *t* | *U* | *p* | *t* | *U* | *p* | *t* | *U* | *p* |
| G1<br>Schaffer | $\bar{X}$ | - | 65 | .142 | 4.4 | - | <.001 | - | 164 | .002 | - | 103 | .821 | -7.2 | - | <.001 |
| | $\sigma$ | - | 71 | .235 | - | 2 | <.001 | - | 112 | .525 | -0.2 | - | .862 | -2.9 | - | .007 |
| | $\kappa$ | - | 99 | .964 | - | 54 | .046 | - | 55 | .052 | 0.8 | - | .419 | 1.8 | - | .079 |
|  | Sk | 0.2 | - | .87 | - | 23 | <.001 | - | 53 | .041 | 0.3 | - | .739 | - | 61 | .098 |
| G2<br>Lac-Mol | $\bar{X}$ | -3.4 | - | .002 | 11.8 | - | <.001 | - | 183 | <.001 | - | 46 | .017 | - | 192 | <.001 |
| | $\sigma$ | -6.7 | - | <.001 | - | 0 | <.001 | -3.1 | - | 0.05 | - | 19 | <.001 | - | 118 | .363 |
| | $\kappa$ | -4.5 | - | <.001 | - | 90 | .751 | -0.1 | - | .962 | 4.5 | - | <.001 | - | 61 | .098 |
|  | Sk | - | 185 | <.001 | 0.2 | - | .83 | 1.6 | - | .113 | - | 9 | <.001 | 2.3 | - | .033 |
| G3<br>PP | $\bar{X}$ | -4.8 | - | <.001 | 3.1 | - | .005 | - | 165 | .001 | - | - | - | -4.7 | - | <.001 |
| | $\sigma$ | -4.8 | - | <.001 | 3.7 | - | .001 | -0.9 | - | .36 | -2.7 | - | .013 | -1.8 | - | .079 |
| | $\kappa$ | - | 113 | .496 | 1.3 | - | .224 | -1.1 | - | .262 | - | 70 | .217 | -1 | - | .328 |
|  | Sk | - | 120 | .316 | - | 34 | .003 | 0.5 | - | .601 | - | 78 | .387 | 0.1 | - | .964 |
| G4<br>GCsom | $\bar{X}$ | -5.4 | - | <.001 | - | 62 | .108 | -5.3 | - | <.001 | -3.4 | - | .002 | -4.9 | - | <.001 |
| | $\sigma$ | -7.4 | - | <.001 | - | 48 | .022 | - | 172 | <.001 | -3.5 | - | .002 | -3.1 | - | .005 |
| | $\kappa$ | -3.7 | - | .001 | - | 70 | .217 | - | 88 | .683 | - | 64 | .13 | 3.6 | - | .001 |
|  | Sk | -4.7 | - | <.001 | - | 62 | .108 | - | 89 | .717 | - | 76 | .339 | 4.7 | - | .001 |
| G5<br>Rem | $\bar{X}$ | - | 170 | <.001 | - | 74 | .294 | - | 177 | <.001 | 3.8 | - | <.001 | - | 193 | <.001 |
| | $\sigma$ | - | 147 | .022 | - | 67 | .170 | -5.1 | - | <.001 | 1.1 | - | .258 | - | 195 | <.001 |
| | $\kappa$ | 2.5 | - | .021 | - | 89 | .717 | - | 170 | <.001 | - | 137 | .072 | - | 184 | <.001 |
|  | Sk | 4.1 | - | <.001 | - | 81 | .467 | - | 167 | <.001 | - | 153 | .01 | - | 178 | <.001 |

